# Termination of DNA replication at Tus-*ter* barriers results in under-replication of template DNA

**DOI:** 10.1101/2021.02.25.432933

**Authors:** Katie H. Jameson, Christian J. Rudolph, Michelle Hawkins

## Abstract

The complete and accurate duplication of genomic information is vital to maintain genome stability in all domains of life. In *Escherichia coli*, replication termination, the final stage of the duplication process, is confined to the ‘replication fork trap’ region by multiple unidirectional fork barriers formed by the binding of Tus protein to genomic *ter* sites. Termination typically occurs away from Tus-*ter* complexes, but they become part of the fork fusion process when a delay to one replisome allows the second to travel more than halfway around the chromosome. In this instance, replisome progression is blocked at the non-permissive interface of Tus-*ter* and termination occurs when a converging replisome meets the non-permissive interface. To investigate the consequences of replication fork fusion at Tus-*ter* complexes, we established a plasmid-based replication system where we could mimic the termination process at Tus-*ter in vitro*. We developed a termination mapping assay to measure leading strand replication fork progression and demonstrate that the DNA template is under-replicated by 15-24 bases when replication forks fuse at Tus-*ter* complexes. This gap could not be closed by the inclusion of lagging strand processing enzymes as well as several helicases that promote DNA replication. Our results indicate that accurate fork fusion at Tus-*ter* barriers requires further enzymatic processing, highlighting large gaps that still exist in our understanding of the final stages of chromosome duplication and the evolutionary advantage of having a replication fork trap.

## INTRODUCTION

In all domains of life, DNA replication is initiated at origins which direct the assembly of multi-subunit replication complexes called ‘replisomes’. Once fully assembled, replisomes travel along the DNA as replication fork complexes which move away from each other in opposite directions. They progress until they reach a chromosome end or converge with a replication fork complex travelling in the opposite direction resulting in a fusion event (1). The number of fusion events is directly correlated to the number of origins in each organism. This means that the number of expected fork fusions varies significantly between organisms; from thousands in metazoa to a few in some archaea and only one in most bacteria (2). The ability to carry out the convergence and fusion of replication fork complexes with high accuracy is essential to the maintenance of genomic stability and cell survival (3). Recent studies in both prokaryotes and eukaryotes have highlighted the fact that the fusion of replication fork complexes requires careful orchestration by a variety of protein co-factors (1, 3–7).

Like the majority of bacteria, *E. coli* has a single circular chromosome which is replicated bidirectionally from a single replication origin (*oriC*). The single origin dictates that each chromosomal half or ‘replichore ‘is duplicated by a single replisome and the number of replication fork fusions is restricted to exactly one, which takes place midway around the chromosome (8) (Figure 1A). The location of termination is constrained by a specialised replication fork trap formed by a series of polar blocks which allow replisomes to enter but not leave the termination region. These polar blocks are created by asymmetric binding of the Tus terminator protein to a series of 23 bp non-palindromic *ter* sequences (*terA–J*) distributed at either side of the termination region (9, 10) (Figure 1A and 1B). Five *ter* sites flank both sides of the termination region. Each replisome is able to bypass the first five sites it encounters in the permissive direction by displacing Tus protein. However, the replisome will be arrested at any Tus-*ter* complexes encountered in the non-permissive orientation (6, 11).

**Figure 1.**
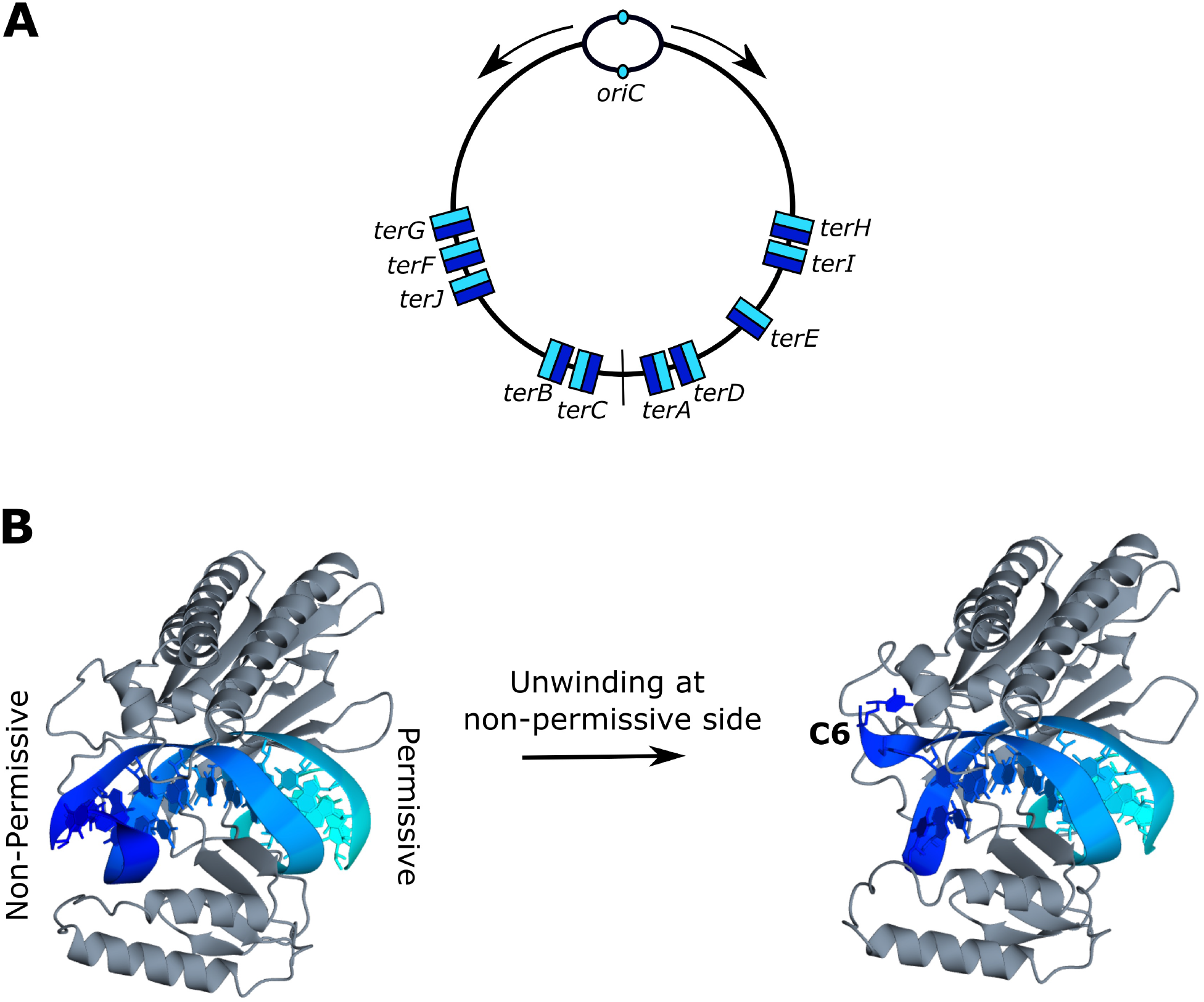
DNA replication and Tus-*ter* termination trap in *E. coli*. **(A)** *E. coli* contains a single circular chromosome which replicates bidirectionally from a single origin (small oval). Direction of replisome travel from the origin is depicted by arrows. Chromosomal midpoint indicated by a straight line. Location of *ter* sites on the *E. coli* chromosome are shown relative to *oriC*. Permissive orientation displayed in light blue, non-permissive orientation displayed in dark blue. **(B)** Structure of Tus-*ter* (PDB ID: 2I06) illustrating the non-permissive and permissive faces (left) and the ‘locked’ conformation formed by DNA unwinding at the non-permissive face (right). The cytosine base at position six of *ter* (C6), which flips into a specific binding site on the non-permissive face of Tus to form the ‘lock’, is indicated.

The asymmetry of replisome arrest at the Tus-*ter* complex has been extensively studied (5, 11). These studies have demonstrated that polar arrest at Tus is triggered by the approaching replisome unwinding the double-stranded DNA immediately adjacent to the non-permissive face (12). This induces specific DNA-Tus contacts which generate a ‘locked ‘complex that cannot be bypassed by the oncoming replication machinery. Specifically, a base on the leading strand template (C6) flips into a specialised binding site on Tus (Figure 1B), guided by nearby residues, to create a sustained barrier (12, 13). Unwinding alone has been shown to be enough to generate a ‘locked’ Tus-*ter* complex (13, 14). Nevertheless, the block may be further stabilised by an interaction between Tus and the replicative helicase, DnaB, which is thought to sit at the head of the replisome (15).

Although the trap system limits replication fork fusions to the termination region, fusions typically occur away from Tus-*ter* sites in the majority of cells. This was demonstrated by early labelling experiments (16) and, more recently, marker frequency analysis produced by Deep Sequencing has shown that fusion typically occurs close to the numerical midpoint of the chromosome (17). Deletion of *tus* from wild type cells results in only mild distortion of nucleic acid metabolism and does not significantly change the location of replication fork fusion, demonstrating that it is not a requirement for replication termination (18, 19). Recent analysis has illustrated that replication fork traps are not widely distributed amongst bacteria (20). However, fork traps are found not only in some Gram-negative bacteria such as *E. coli*, but also in the Gram-positive *Bacillus subtilis*. These two fork trap systems show no significant sequence or structural similarity, indicating that they have evolved via convergent evolution (11). This indeed suggests that the fork trap conducts an important physiological function.

Multiple functions of the fork trap have been proposed. One early suggestion was the potential to contain over-replication of the genome (21). This theory has been bolstered in recent years by the observation of over-replication of the terminus region in the absence of RecG (17) and 3’-exonucleases (22, 23). These enzymes are thought to be involved in the processing of intermediates that form as a result of fork fusion events and can lead to replication restart if the intermediates persist for longer than normal (8, 23). This model is supported by the fact that over-replication is either eradicated or significantly reduced in cells lacking either RecG or 3’-exonucleases following linearisation of the chromosome in the termination area, which will prevent replisomes from fusing (23–25). Additionally, over-replication in the absence of RecG can be induced in ectopic chromosomal locations if replisomes are forced to fuse in these ectopic locations (26). These observations are in line with the idea that intermediates formed as part of termination can trigger replication restart and lead to over-replication of the chromosome (17, 23). Thus, one important role of the fork trap might be to block forks initiated within the termination area travelling towards the origin.

Another scenario where the replication fork trap will come into play is when one replisome is stalled before it reaches the chromosomal midpoint. Studies with ectopic chromosomal origins suggest that Tus-*ter* is an effective barrier *in vivo* when one replisome is delayed and it has been shown that a subset of fusion events occur at Tus-*ter* barriers *in vivo* (27). In this scenario, fusion must occur at Tus-*ter* when a replisome paused at the non-permissive face of Tus is met by a moving replication fork complex coming towards the permissive face. Surprisingly little is known about what happens in this scenario as biochemical studies of Tus-*ter* have typically focused on understanding what happens when a replisome approaches Tus-*ter* from either the blocking or permissive orientation, but not when replisomes meet at Tus-*ter*.

A recent *in vitro* study found that the addition of Tus-*ter* to a plasmid replication system at a location mimicking that of the normal chromosomal arrangement, inhibited the formation of circular monomers (28). This inhibitory effect could be overcome by increasing the spacing between Tus-*ter* such that it was likely not to be the fusion point of the replication forks; or by adding UvrD (28), an accessory helicase reported to overcome Tus-*ter* barriers (29). This result suggests that additional processing steps are required for the fusion of a replisome paused at Tus-*ter* with a freely moving replication fork complex.

We have designed an *in vitro* DNA replication system where we can control the fusion of two replication forks at a Tus-*ter* complex and monitor how far each replisome progresses during replication fork fusion. Our data demonstrate that the fusion of a fork paused at a Tus-*ter* complex with a freely moving fork will not result in fully replicated templates. This fusion scenario leaves a gap of 15–24 bp of DNA between the two replisomes that remains unreplicated. These results raise further questions about how fork fusion events are completed at Tus-*ter in vivo* and renew the debate about the physiological advantage of having a replication fork trap.

## MATERIAL AND METHODS

### Plasmids

Backbone fragments of pKJ1 were purchased from ThermoFisher Genestrings service. Two dsDNA fragments of 2269 and 2945 bp were synthesised based on pIK02 (30). The fragments were designed to retain the *E. coli* replication origin, *oriC*, the plasmid initiation site *colE1*, the Tus DNA-binding site, *terB*, and the ampicillin resistance gene, *bla*. Recognition sequences for four unique ssDNA nicking enzymes were included, two approximately 100 bp upstream and two approximately 100 bp downstream of *terB* (Figure 3A). All other instances of these ssDNA nicking sequences were removed in the design of the synthetic fragments. A repeat region of DNA containing 22 *lacO* binding sites was amplified from pIK02 by PCR using the following primers: 5’-GCCAGCACGTAGCTAGCAAACCG-3’ and 5’-CCTTCTAGAGATTCGACTCTAGAGTCC-3’. Fragments were annealed by ligation independent cloning using T4 polymerase (NEB) to generate complementary 5’-overhangs, as described (31).

### Protein Production

Proteins used in replication assays were purified as previously described (32). Rep and UvrD were purified as described (33). RecG was a kind gift from Robert Lloyd and Geoff Briggs. RNase HI, DNA ligase and DNA polymerase I encoding genes were cloned into a modified pET28a vector which encodes an N-terminal His_6_SUMO-tag. Proteins were overexpressed in BL21 (DE3). Cells were grown at 37 °C to an OD_600_ of 0.6–0.8 before expression was induced by the addition of 1 mM IPTG. Growth was continued at 18 °C for approx. 18 hours before cells were harvested by centrifugation and resuspended in a buffer of 50 mM Tris pH 8.0, 500 mM NaCl, 1 mM DTT and 20 mM imidazole. Cells were lysed by sonication and cell debris pelleted by centrifugation. DNA was precipitated from soluble cell lysate by the dropwise addition of Polyethylenimine to a final concentration of 0.075 % (v/v) and stirring for 20 minutes at 4 °C. Precipitated DNA was pelleted by centrifugation and soluble lysate was loaded onto a 5 ml HisTrap HP column (GE Healthcare). The column was washed with 7 column volumes of loading buffer before being eluted with a gradient of imidazole (20–500 mM) over 20 column volumes (100 ml). Fractions corresponding to the presence of the protein of interest were combined according to chromatographic analysis at *A280* and SDS-PAGE analysis. The His_6_SUMO tag was cleaved by the addition of ULP1 and incubation overnight at 4 °C. The protein was simultaneously dialysed into 50 mM Tris pH 8.0, 500 mM NaCl, 1 mM DTT to remove imidazole from the buffer. Cleavage products were passed over a second HisTrap HP column, which was washed and eluted in an analogous manner to the first. Flow-through fractions containing the protein of interest were combined and concentrated to <1 ml using a centrifugal concentrator. Proteins were further purified by gel filtration. RNase HI was loaded onto a 16/600 Superdex 75 column (GE Healthcare), whilst PolI and Ligase were further purified on a 16/600 Superdex 200 column (GE healthcare). Following gel filtration, Ligase was additionally dialysed into 50 mM Tris pH 7.5, 75 mM NaCl, 1 mM DTT and loaded onto a 1 ml HiTrap Q column and eluted with a gradient of salt (75–1000 mM NaCl) over 30 column volumes. RNase HI, Ligase and PolI were dialysed into a storage buffer of 50 mM Tris pH 7.5, 100 mM NaCl, 2 mM DTT, 40% glycerol before storage at −80 °C. Tests were carried out to confirm protein activities in replication assay buffer conditions. No protein shows significant dsDNA nuclease activity over the timeframe of the assays used in this study.

### Replication Assay

Replication reactions were carried out in 40 mM HEPES pH 8.0, 10 mM Magnesium Acetate, 10 mM DTT, 2 mM ATP, 200 µM GTP/UTP/CTP, 40 µM dNTPs and 0.1 mg/ml BSA. Each reaction (15 µl final volume) contained 200 ng template DNA (approx. 2 nM), 50 nM polymerase III core (αεθ), 25 nM clamp loader complex (τ_3_δδ’χψ), 160 nM DnaB, 160 nM DnaC, 1 µM SSB, 80 nM β-clamp, 30 nM HU, 200 nM DnaG and 133 nM Gyrase (A_2_B_2_). Where indicated, reaction mixes contain 400 nM Tus and/or 400 nM LacI. Reaction mixes were assembled on ice, then incubated at 37 °C for 2 mins before replication was initiated by addition of 300 nM DnaA. Reactions were then incubated at 37 °C for 2 min before the addition of 30 units of SmaI (Promega), 46 kBq [a-^32^P]-dCTP (111 TBq/mmol) and, where indicated, 1 mM IPTG. Reactions were incubated for a further 2 min at 37 °C before being terminated by the addition of 2.5 µl STOP buffer (2.5% SDS, 200mM EDTA and 10 mg/ml proteinase K). Reaction products were precipitated by ethanol precipitation and resuspended in 25 µl 50 mM NaOH, 30 mM EDTA before analysis by denaturing gel electrophoresis (0.7 % agarose in 30 mM NaOH, 2mM EDTA). 5′-labelled HindIII-digested λ DNA was used as a marker. Samples were run on a 40 cm gel for 400 Vh (typically 16 h at 25 V) before being dried and analysed by phosphorimaging and autoradiography. One-way assays were carried out as above except for the exclusion of Gyrase and SmaI was substituted for 30 units Ncol-HF (NEB) or 30 units SacI-HF (NEB). Reactions containing RLP include 4 nM RNase HI, 25 or 50 nM ligase and 30 nM DNA polymerase I. Where indicated, Rep, UvrD or RecG were included at a concentration of 200 nM.

### Termination Mapping

Replication reactions to map replication end points were carried out as described above with a 4 × reaction mix (60 µl). Following ethanol precipitation, reaction products were resuspended in 50 µl relevant manufacturer’s enzyme buffer (1 × Cutsmart (NEB) for nicking with Nt.BspQI or 1 × Buffer R (ThermoFisher) for nicking with Nb.Bpu101). Products were subsequently split into two 25 µl aliquots; one of which was incubated with 1 µl nicking enzyme (10 units Nt.BspQI (NEB) or 5 units Nb.Bpu101 (ThermoFisher)) and the other, 1 µl enzyme buffer (control). Reactions were incubated at 37 °C for 30 min then at 80°C for 20 min. Products were split into two 12.5 µl samples. To one sample, 6 µl STOP solution (USB Thermo Sequenase Cycle Sequencing Kit, Affymetrix) was added for analysis by Urea PAGE gel electrophoresis. The other sample was precipitated by ethanol precipitation and resuspended in 12.5 µl 50 mM NaOH, 30 mM EDTA for analysis by denaturing gel electrophoresis (see *replication assay* above).

Cycle sequencing reactions were carried out according to the manufacturer’s instructions (USB Thermo Sequenase Cycle Sequencing Kit, Affymetrix) with minor modifications. In brief, primers were 3’-labelled with [a-32P]-dCTP in reaction mixes containing dGTP dATP but excluding dTTP. 3’-labelling was carried out using thermocycling between 95 °C (15 s) and 60 °C (30s) for 50 cycles. The termination step was also carried out in the presence of all four dNTPs and dd-NTPs for chain termination. During the termination step, reactions were thermocycled between 95 °C (30 s) and 72 °C (90 s) for 50 cycles. The leading strand approaching the Tus-*terB* non-permissive face (clockwise travelling replisome) was sequenced with the primer 5’-AATGCTTATTATCATGACATTACCTATCC-3’ whist the leading strand approaching Tus-*terB* permissive face (counter-clockwise travelling replisome) was sequenced with the primer 5’-TAAGGCATTTGTTTCAGGTTACTCC-3’. Nicked reaction products and sequencing products were analysed alongside one another by 8 % Urea TBE PAGE. Gels were run in 1% TBE at 50 W for 120 min. Gels were dried and analysed by phosphorimaging and autoradiography.

## RESULTS

### Controlled fusion of replisomes at Tus-*ter*

We used an *in vitro* plasmid replication system to study what happens when a moving replication fork complex fuses with another replication fork complex paused at Tus-*ter*. In the presence of DNA gyrase and reconstituted *E. coli* replication machinery, bidirectional DNA replication can be initiated from an *E. coli oriC* sequence on a plasmid template *in vitro* (34). Replisomes are able to progress around the majority of the template but eventually stall due to positive supercoiling, leaving the last ∼150 bp of DNA unreplicated. This tension can be released by linearisation of the plasmid template with a restriction enzyme, thereby allowing replication of the last 150 bp of DNA (34). Here, we initiated replication from the *oriC*-containing plasmid, pKJ1, which can be linearised by the addition of SmaI (Figure 2A). Unimpeded, the major product of this reaction is a 6 kb band which corresponds to full-length leading strand replication products from one of the two replisomes progressing around the entire template before the release of the other (Figure 2B, lane 1). The processive nature of the replisome means that *E. coli* can replicate DNA at approx. 600-800 bp/sec (35, 36), thus if there is a >10 sec delay in the release of one replisome from *oriC*, the other replisome is able to complete replication before the second replisome has been released. A smear of products is also observed at <6 kb corresponding to leading strand products from replisomes meeting stochastically throughout the plasmid. Lagging strand products are observed as a smear of products centred around 0.5 kb (Figure 2B, lane 1). Replisome progression can be blocked by the use of protein binding sites on the template DNA. We have taken advantage of two such blocks to generate a system where we are able to block replisomes travelling in both directions, before releasing one to allow fusion to occur in a controlled manner.

**Figure 2.**
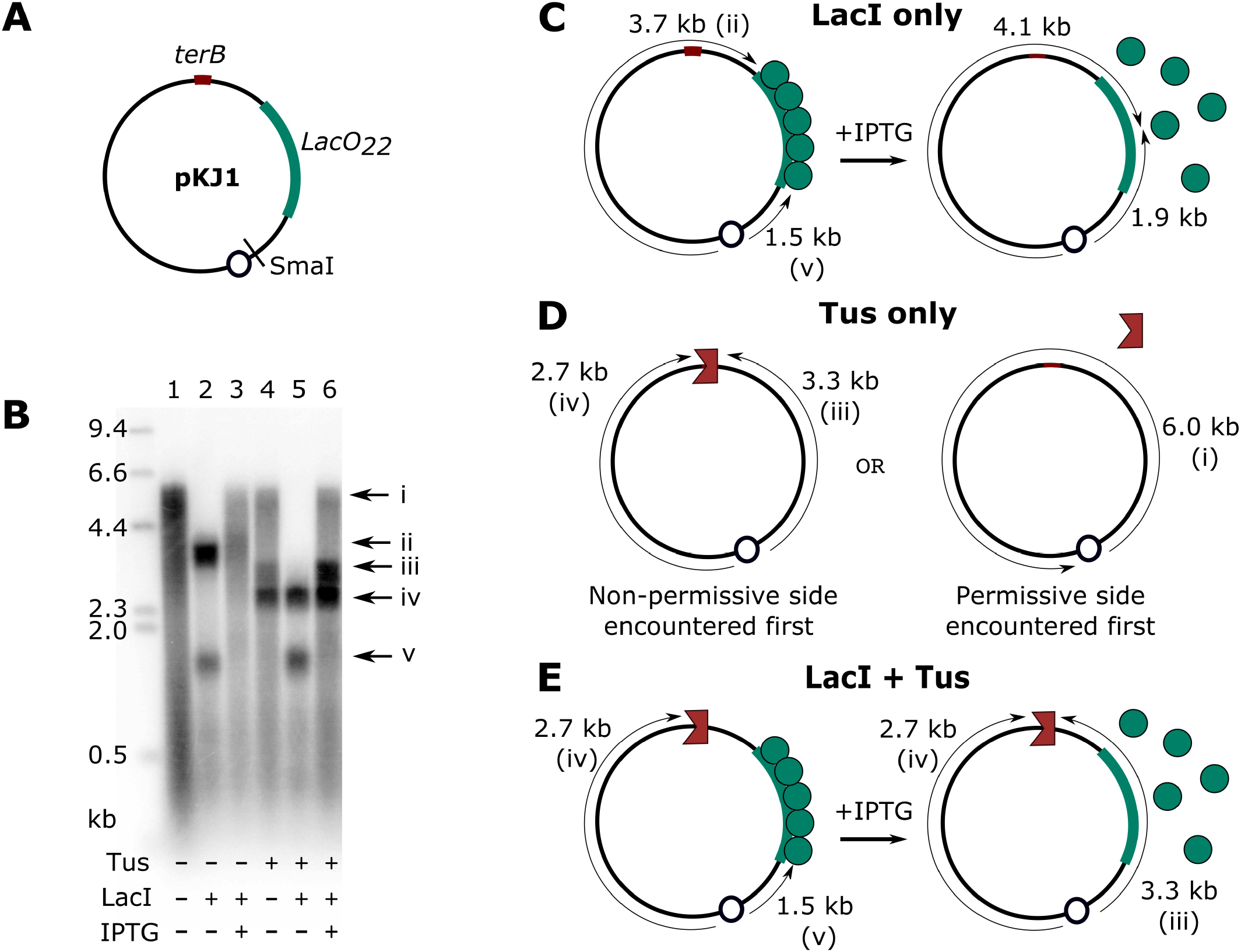
Replication fork fusion can be controlled to occur at Tus-*terB*. **(A)** pKJ1 replication assay template, indicating the location of replication initiation at *oriC* (small circle), directionality of replisome travel and the SmaI cleavage site. The locations of *lacO*_*22*_ and *terB* are indicated. **(B)** Denaturing agarose gel of replication products from pKJ1, in the presence and absence of LacI, Tus or both proteins, and with and without addition of IPTG, as indicated. Band labels correspond to replication products shown in panels C-E. **(C)** Replisome movement in the presence of LacI and its release by the addition of IPTG, with expected leading strand product lengths. **(D)** Replisome movement in the presence of Tus, including expected leading strand replication products, depending on which side of Tus is encountered first. **(E)** Replisome movement in the presence of LacI and Tus, before and after release of LacI by addition of IPTG. Expected leading strand replication product lengths are indicated.

Replisomes can be blocked at a *lacO*_*22*_ array bound by LacI and subsequently released by the addition of IPTG (30), enabling them to continue replication (Figure 2C). Replisomes can also be blocked at Tus bound *ter* sequences if the replisome encounters Tus-*ter* in the non-permissive direction (Figure 2D). Replisomes that encounter Tus-*ter* in the permissive direction are able to displace Tus and travel all the way round the template (37) (Figure 2D). Unlike LacI-*lacO* blocks, replisomes blocked at Tus-*ter* cannot be released. We have used both blocks to create a system where we can control the fusion of two replication forks directly at a Tus-*ter* complex. We initiate bidirectional replication on pKJ1, block replisomes travelling in the clockwise direction at Tus-*terB*, and those travelling in the counter-clockwise direction at *lacO*_*22*_ (Figure 2E); this generates leading strand products of 2.7 kb and 1.5 kb, respectively (Figure 2B, lane 5). We then release the replisome blocked at *lacO*_*22*_ by addition of IPTG, allowing it to resume synthesis and consequently meet the blocked replisome at Tus-*terB*, yielding 2.7 kb and 3.3 kb products (Figure 2B, lane 6 and 2E). A small quantity of full-length product (6 kb) is also always observed in these reactions. We reason that this is likely due to a small percentage of template molecules which have replisomes travelling unidirectionally counter-clockwise and hit Tus-*terB* from the permissive side (Figure 2D). We anticipate a similar number of unidirectional replisomes travelling in the clockwise direction, but these cannot be quantified because they contribute to the 2.7 kb products.

We confirmed that both blocks work independently of one another using the same template and reaction conditions. As expected, replisomes blocked at a *lacO*_*22*_ array bound to LacI generate products of 3.7 kb (clockwise) and 1.5 kb (counter-clockwise) (Figure 2B, lane 2 and 2C). Release by the addition of IPTG produces a smear of products from replisomes meeting within the *lacO* array (3.7–4.1 kb) and (1.5–1.9 kb) and full-length (6 kb) products (Figure 2B, lane 3). The addition of Tus alone produces a mixture of products from replisomes which are blocked and meet at Tus-*terB* (2.7 kb, clockwise, 3.3 kb, counter-clockwise), and 6.0 kb full length products which first reach Tus-*terB* from the permissive side (counter-clockwise) (Figure 2B, lane 4 and 1D).

### Replisomes fusing at Tus-*ter* leave a 15–24 bp gap

Using the replication assay described above, we analysed the length of the leading strand products from each replisome to understand how far each replisome can travel during fusion at Tus-*terB*. In an assay we call ‘termination mapping’, we introduced unique single strand nicking sites approximately 100 bp upstream of *terB* on each strand of the replication template and used these to cut out individual leading strands from the final replication products (Figure 3A). We analysed the size of these products to determine the stop sites of the leading strand polymerases during the reaction. Nicked products were analysed by denaturing urea PAGE, with reference to sequencing products from cycle sequencing reactions carried out using the same template DNA. The DNA binding location of the 5’-teminus of the primers used for cycle sequencing was identical to the nicking site (Figure 3A), meaning that comparison of the length of the nicked product with the sequencing reaction products provided a direct read out of the 3’-stop site of DNA replication. Nicked products (N, Figure 3B and 3C) were compared with products from the same reaction that had not been treated by a nicking enzyme (C, Figure 3B and 3C), to ensure there was no misinterpretation of products generated during the replication reaction. Any products observed across both nicked and untreated lanes were ignored. This analysis showed that synthesis of DNA from replisomes travelling clockwise towards Tus-*terB* in the non-permissive direction, stopped at an adenosine immediately upstream of *terB* (Figure 3B and 3D). This is in line with previous replisome mapping studies which found that the main leading strand stop location at the Tus-*ter* non-permissive interface one base closer to *ter* (37, 38). Leading strand synthesis in the opposing direction, travelling towards Tus-*terB* in the permissive direction, stopped at several locations immediately upstream of and within *terB* (Figure 3C and 3D).

**Figure 3.**
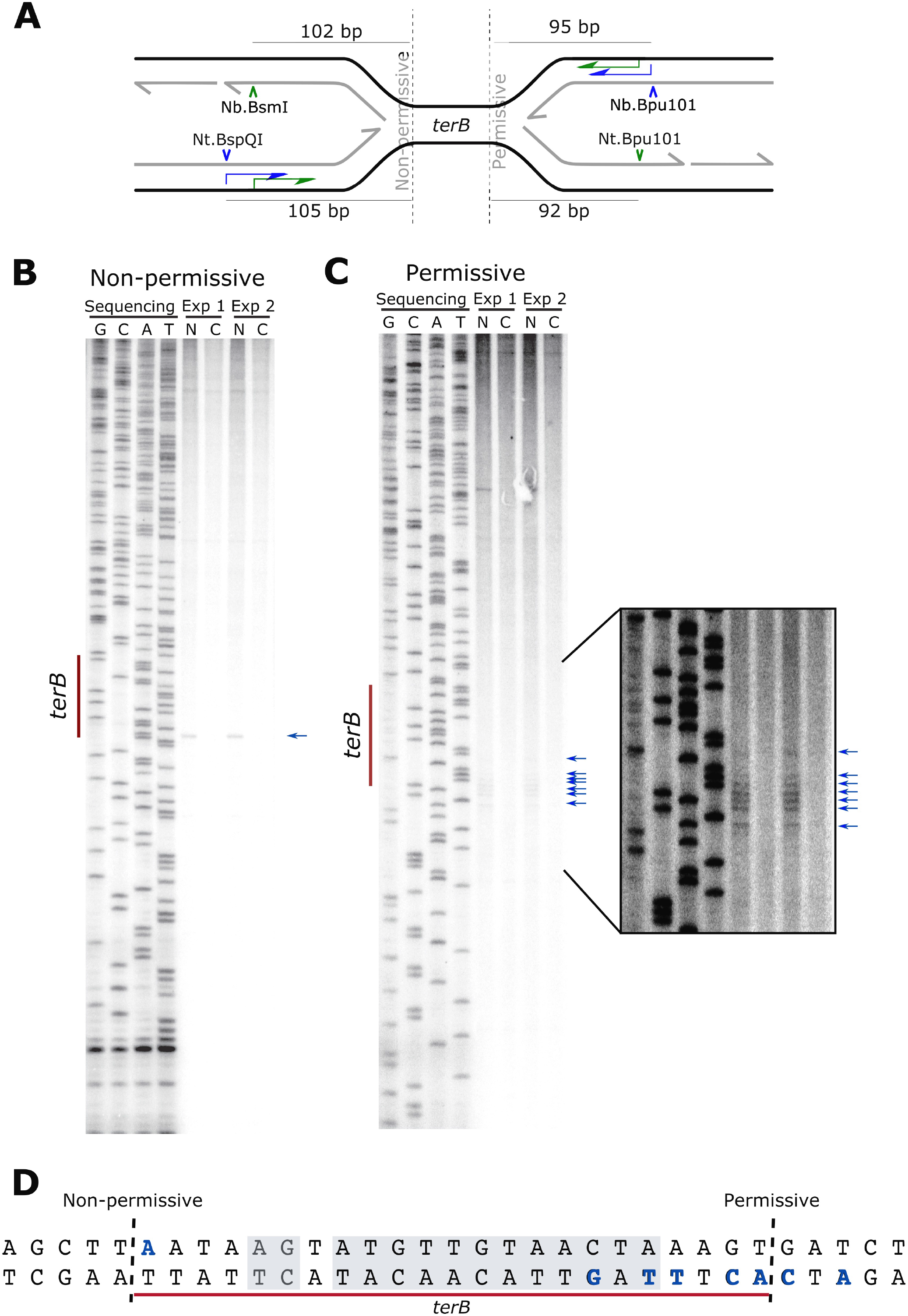
Replication fork complexes meeting at Tus-*ter* leave a gap of 15-24 bp of unreplicated DNA. Analysis of leading strand products created during replisome fusion at Tus-*terB* **(A)** Locations of single strand nicking sites and primer annealing sites for cycle sequencing on pKJ1, with respect to *terB*. The 5’ end of each primer corresponds to a single nicking site, enabling replication product stop sites to be determined. **(B)** Mapping analysis of the leading strand product approaching Tus-*terB* in the non-permissive direction. Nicked products (stop sites) are indicated by arrows. **(C)** Mapping analysis of the leading strand product approaching Tus-*terB* in the permissive direction. Nicked products (stop sites) are indicated by arrows. **(D)** *terB* sequence indicating leading strand stop locations (bold) and the Tus binding site (shaded area).

We attempted a similar analysis of the lagging strand products, this time introducing a nicking site on the strand complementary to the sequencing products (Figure 3A). The nicking site corresponded to the 5’ end of the primer (but on the complementary strand). Comparison of nicked and cycle sequencing products would therefore allow us to identify the start sites of lagging strand synthesis. Again, nicking sites were introduced approximately 100 bp upstream and downstream of *terB*. However, we observed no products specific to our nicking enzyme analysis. Reasoning that we may be too close to *terB* to observe a lagging strand start site, we moved the nicking sites approximately 1000 bp away from *terB* and performed a wider analysis. Again, we observed no products which could be clearly identified as specific to the nicking analysis. We suspect therefore that the lagging strand start sites are too stochastic to be observed by our mapping method.

Although we cannot map the precise location of the lagging strand start sites nearest to Tus-*terB*, the stop locations of the leading strands provide an insight into how far each replisome is able to travel during fork fusion at Tus-*terB*. Taking both the minimum and maximum distances between the ends of both leading strand products, a gap of 15–24 bp of DNA remains unreplicated during fusion at Tus-*terB*.

### Unreplicated DNA at Tus-*terB* is a consequence of fusing replisomes

To confirm that the gap of unreplicated DNA observed in our mapping analysis is a direct consequence of two replisomes fusing at Tus-*terB*, we analysed what happens when replisomes are limited to travel in only one direction towards Tus-*terB* (i.e. approaching the permissive or non-permissive face). To control the direction of travel of the replisome through Tus-*terB*, we carried out experiments in the absence of gyrase. In these reactions, only one replisome is able to travel to any extent away from *oriC*, replicating approximately 1 kb of DNA before pausing due to the accumulation of positive supercoils in the DNA. This replisome can be subsequently released by linearisation of the template with a restriction enzyme. As long as the restriction enzyme cleaves the template behind the progressing fork, it can then continue replicating the remainder of the DNA template (34). Replisome release is stochastic, meaning an equal number of replisomes are released in either direction (21). When we linearise with SmaI (Figure 4A i), a replication fork which has progressed in either direction will be released. We have taken advantage of this phenomenon by introducing template linearisation sites > 1 kb away from *oriC* (Figure 4A i); in these reactions we cut the template in front of one fork, removing the template in one direction. In this way we are able to analyse only products from replisomes travelling towards Tus-*terB* in either a clockwise or counter-clockwise direction. Thus, by linearising the template at NcoI (1.9 kb clockwise), all clockwise (non-permissive) products can only progress to 1.9 kb in length, and we can analyse what happens when replisomes travel towards Tus-*terB* in the permissive direction without a replisome blocked at the non-permissive side (Figure 4A, iii). Conversely, if we cut with SacI (1.6 kb counter-clockwise), all counter-clockwise replisomes can only replicate 1.6 kb of DNA, and we can analyse what happens to replisomes approaching Tus-*terB* in the non-permissive direction (Figure 4A, ii).

**Figure 4.**
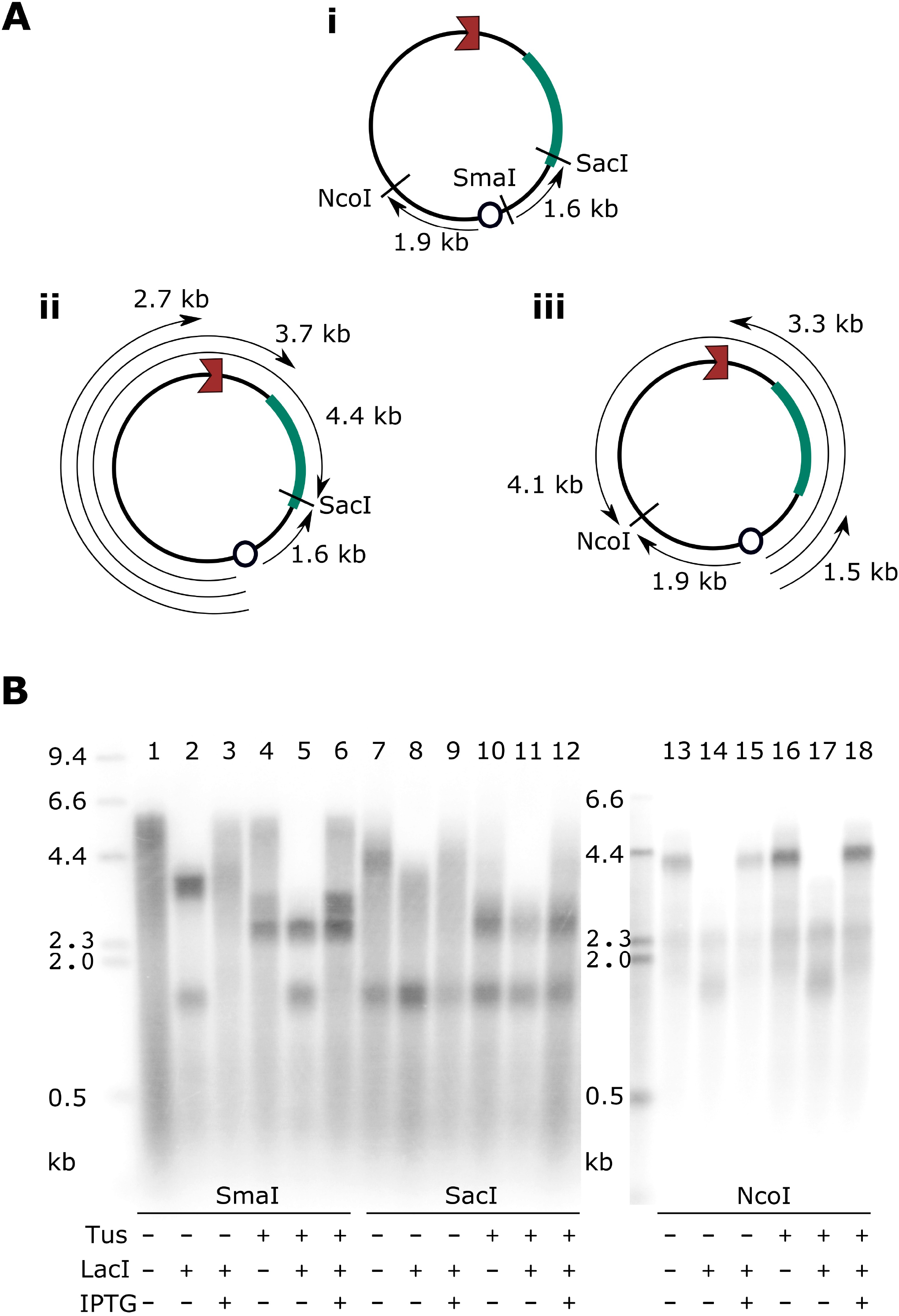
The unreplicated gap at *terB* is a direct consequence of replisomes meeting at Tus-*ter*. Analysis of unidirectional replisome travel towards Tus-*terB* **(A)** (i) Locations of SmaI, NcoI and SacI, with respect to the replication origin. (ii) Hypothetical leading strand products if the template is linearised by SacI cleavage (iii) Hypothetical leading strand products if the template is linearised by NcoI cleavage **(B)** Denaturing agarose gel of replication products linearised at SmaI, SacI or NcoI as indicated.

When templates are linearised with SacI (analysing replisomes travelling clockwise towards Tus-*ter* non-permissive face), we consistently see the expected 1.6 kb product from counter-clockwise travelling replisomes. In the presence of Tus, we see products corresponding to replisomes that have been blocked at Tus-*terB* in the clockwise direction (2.7 kb band, Figure 4A, ii and 4B, lanes 10–12), indicating that Tus blocks replisomes in this linearised system.

When templates are linearised with NcoI (analysing replisomes travelling towards Tus-*terB* permissive face), products are seen which correspond to replisomes travelling through Tus-*terB* in both the presence and absence of Tus (4.1 kb bands, Figure 4A, iii and 4B, lanes 13 and 16), and after the release of the counter-clockwise replisome from *lacO*_*22*_ (4.1 kb band, Figure 3B, lane 18). These data confirm that the halting of replisome travel in the counter-clockwise (permissive) direction (3.3 kb band, Figure 2D, lane 6 and Figure 4B, lane 6) requires the presence of a replisome at the non-permissive face on the other side of Tus. This confirms that the unreplicated DNA region seen at Tus-*terB* in our termination mapping assay is a direct consequence of a replisome fusion event at Tus-*terB*.

### The replication gap cannot be closed by the lagging strand processing enzymes, RNase HI, DNA Ligase and DNA polymerase I

Our standard replication reactions lack RNase HI, DNA Ligase and DNA polymerase I (called RLP from here onwards for brevity), enzymes required for joining lagging strand fragments (and by extension, fusing lagging and leading strands to one another). Thus, we investigated whether the addition of these enzymes to our replication assay could process the observed gap in replication. We reasoned that closing the gap would result in an increased proportion of full-length replication products compared to all other products from the same reaction; therefore we compared the percentage of full-length replication products in the presence and absence of RLP. Although we consistently observed an increased intensity across all replication products and a small increase in the percentage of full-length products (<10%) there was not the substantial increase in the proportion of full-length product we would expect if RLP was able to promote complete termination of DNA replication at Tus-*ter* (Figure 5A, compare lane 1 to lanes 5 and 6). This indicates that the lagging strand processing enzymes are insufficient to resolve termination events at Tus-*ter*. Note that an additional band at 4.2 kb is observed in reactions containing RLP due to the ligation of 1.5 kb (LacI-blocked) and 2.7 kb (Tus-blocked) fragments at the origin (Figure 5C, before release from *lacO*_*22*_).

**Figure 5.**
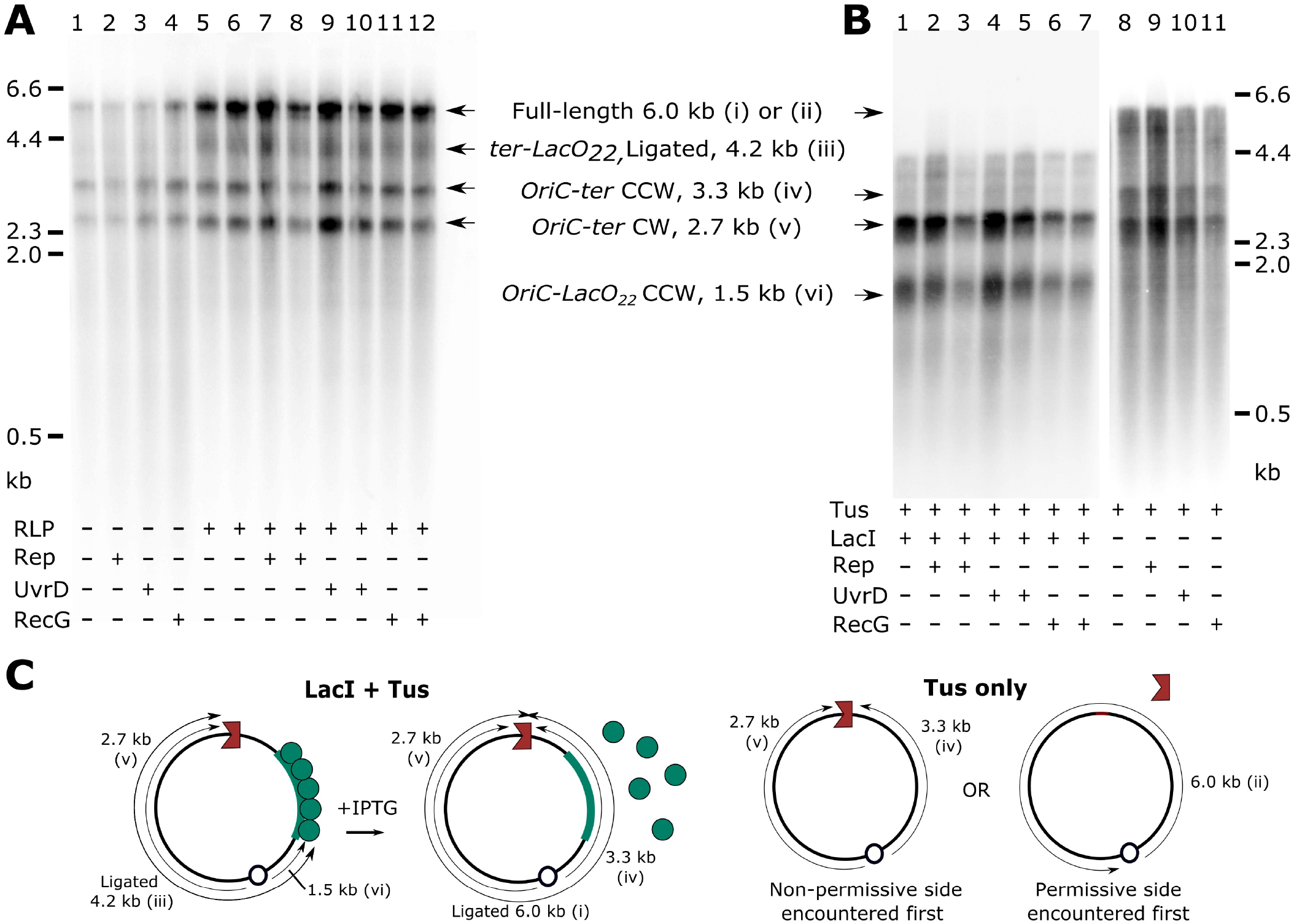
Rep, UvrD or RecG do not assist replication fork fusion at Tus-*ter in vitro*. **(A)** Denaturing agarose gel of replication products from controlled fusion reactions at Tus-*ter*. Addition of RLP in the presence or absence of Rep, UvrD or RecG is unable to increase the proportion of full-length replication products. **(B)** Denaturing agarose gel of replication products blocked at Tus-*ter* and LacI-*lacO*_*22*._ Addition of Rep, UvrD or RecG does not assist replisome progression through Tus-*ter*. **(C)** Predicted replication products from reactions containing Tus and LacI or Tus-only. Replication product labels correspond to labelled bands in panels A-B.

### The replication gap cannot be closed by the accessory helicases Rep and UvrD or by the helicase RecG

Several *E. coli* helicases have been associated with successful DNA replication events *in vivo*. The accessory helicases Rep and UvrD are known to assist replisome progression by removing barriers which would otherwise impede oncoming replisomes (39, 40). Of note, UvrD has previously been implicated in overcoming Tus-*ter* barriers both *in vivo* (29) and *in vitro* (28). In addition, the helicase RecG appears to play a role in the successful termination of DNA replication *in vivo* (17, 26). We therefore hypothesised that the addition of one of these helicases to our assay may facilitate processing of the gap in replication we observed at Tus-*ter*. We first investigated whether Rep, UvrD or RecG could overcome the Tus-*ter* barrier when the replisome approaches Tus-*ter* in a non-permissive direction. We saw no progression beyond Tus-*ter* when replisomes were blocked in both directions (Figure 5B lanes 2–7 and Figure 5C,) and no increase in full-length products compared to a control in the absence of the helicases when replisomes are blocked only by Tus (Figure 5B, compare lane 8 with lanes 9–11, and Figure 5C), indicating that the helicases were unable to assist the replisome in progression through the Tus-*ter* barrier.

We next added Rep, UvrD or RecG to our fusion assay where one replisome is released at *lacO*_*22*_ to meet the other replisome at Tus-*ter*. Although the helicases are unable to overcome the Tus-barrier in the non-permissive direction, we reasoned they may be able to remove Tus in the permissive direction in our full replication assay, facilitating strand fusion. Again, we analysed the proportion of full-length replication products compared to all replication products in the presence of RLP. Addition of UvrD, Rep or RecG produced no increase in observed full-length products (Figure 5A, compare lanes 5 and 6 with 7–12). These results demonstrate that neither UvrD, Rep or RecG are able to facilitate strand fusion at Tus-ter, leaving open the question of how the observed gap is closed in living *E. coli* cells.

## DISCUSSION

Termination of DNA replication in bacteria occurs when two replisomes translocating away from the origin in opposite directions meet in the termination area opposite *oriC*. For a successful termination event to occur, nascent leading and lagging strands need to be ligated, the replisomes disassembled and the resulting fully-replicated chromosomes decatenated by topoisomerases. In *E. coli* the majority of fusion events occur at the chromosomal midpoint away from Tus-*ter* sites, indicating that both replisomes travel with approximately the same speed (16, 17). However, replisomes can get delayed at obstacles such as a nucleoprotein block, DNA secondary structures or DNA lesions. If one fork is significantly delayed, then the second fork will proceed until it is paused at a non-permissive Tus-*ter* face. This fork will remain stalled at Tus-*ter* until the delayed replisome arrives on the permissive side, thereby forcing termination to occur directly at a Tus-*ter* complex. Forks blocked at Tus-*ter* complexes can be easily visualised in normally growing cells, highlighting that they are a regular occurrence (27).

Our characterisation of a fusion event at a Tus-*terB* complex *in vitro* has highlighted an inability for established replisome components to complete replication termination at Tus-*terB* on their own, leaving a gap of at least 15 bp of unreplicated DNA across the *terB* site. Given the similarity of the *E. coli ter* sites and particularly those most commonly used *in vivo*, the innermost sites *terC, terA* and *terB* (27), we believe that our *in vitro* results accurately represent the physiological situation for termination events at Tus-*ter in vivo*. Thus, our results indicate that additional protein activities are necessary to complete DNA replication if one fork is paused at a Tus-*ter* complex and the second fork approaches from the permissive side. Indeed, the analysis of genetic data has led to the hypothesis that a fusion event between two freely moving forks will generate very different intermediates to the situation where one replisome blocked at the non-permissive face of Tus-*ter* fuses with a freely moving replication fork complex that approaches from the opposite direction (23, 26) However, we previously observed that when DNA replication is forced to terminate at Tus-*ter* due to an additional ectopic replication origin, the majority of cells grow without much ill-effect (41, 42). Thus, cells are able to process this type of fusion event without much difficulty.

Two obvious additional requirements for the fusion of replication forks at a Tus-*ter* complex are the displacement of Tus from the *ter* site, and the replication of the *ter* site itself prior to the ligation of leading and lagging strands. The enzymes DNA polymerase I, RNase HI, and DNA ligase are responsible for replacing RNA primers with DNA and joining adjacent lagging strands to one another during DNA replication. As a directly equivalent process, they would also be anticipated to ligate the nascent leading strand with the final lagging strand produced by converging replisomes during the termination of DNA replication. Yet, the addition of PolI, DNA ligase and RNase HI did not result in the gap being closed in our controlled fusion reactions, suggesting that at least one additional processing factor is necessary to allow DNA replication to successfully complete at Tus-*ter*.

It has been established that Tus is removed from *ter* when approached by a replisome (or the replicative helicase) from the permissive direction (12, 13). However, to our knowledge, whether the same applies when another replisome is already blocked at the non-permissive face has not been determined. We observed a marked similarity between the Tus-binding footprint and the unreplicated gap observed in our controlled fusion reactions (Figure 3D). At first glance, the final two leading strand stop sites on the permissive side appear to overlap with the Tus-binding footprint, but closer analysis of the Tus-*ter* locked-structure (12) shows that the leading strand template at these stop sites makes very few contacts with Tus and the majority of these bases are solvent-exposed. In fact, it is the lagging strand template that is much more tightly bound by Tus on the permissive side. This suggests that it is possible for the leading strand template to be available up to the final stop residue without Tus being removed from the DNA. This possibility is especially easy to envisage if the *E. coli* replisome is topologically similar to that of the T7 replisome, where both helicase and polymerase have recently been shown to sit at the head of the replisome, synergistically unwinding the DNA duplex (43).

If the replisome is unable to remove Tus itself, what additional factor is required to displace Tus and allow replication to be completed? It has been previously noted that *E. coli* sometimes appears to be able to overcome Tus-*ter* barriers *in vivo* (44). Likely candidates for this role are the *E. coli* accessory helicases Rep and UvrD. Numerous investigations have been carried out previously to understand if Rep or UvrD are able to displace Tus when travelling towards the non-permissive interface, with contradictory results (45–48). In some experimental conditions, both proteins appear to be blocked by Tus (45) or able to displace Tus (48) when approaching the Tus non-permissive interface, whilst others showed that Rep (46) or UvrD (47) were able to displace Tus. UvrD has been further implicated as a candidate for displacing Tus *in vivo* by a report that the gene is required for viability of a strain carrying ectopic *ter* sites designed to block normal replisome progression (29). A recent *in vitro* study also demonstrated that UvrD could promote the formation of circular replication products which were otherwise absent in the presence of Tus due to the inhibitory orientation of *ter* sites on the template (28). However, when we included these proteins in our reconstituted replisome fusion assays, we did not observe any ability of Rep or UvrD to displace Tus when approaching in the non-permissive direction. This is a striking difference to the ability of Rep and UvrD to displace RNA polymerase and facilitate replisome progression in a directly analogous assay (40). Moreover, Rep and UvrD were unable to promote successful ligation of leading and lagging strands from replisomes converging at Tus-*ter*. These results strongly suggest that they are either not responsible for displacing Tus *in vivo*, or that an additional stimulatory factor is necessary for them to be able to do so.

Another helicase implicated in facilitating fork fusion is RecG. RecG has been shown to play a role in processing intermediates which result from fusion events that otherwise result in replication restart from the terminus region. However, RecG was neither able to displace Tus from the DNA or promote ligation of strands during fusion at Tus-*ter in vitro*. This again suggests that another unknown molecule, or stimulatory factor, is required for successful termination at Tus-*ter*. Indeed, in cells that carry an additional ectopic replication origin, which results in one replication fork complex almost always being blocked at Tus-*ter*, it has been shown that both RecG and Rep can be inactivated without much ill-effect to the cells (26, 40).

Our results pose the obvious question as to the evolutionary advantage of Tus-*ter* for the *E. coli* cell. A recent analysis has shown that sequences related to Tus are found in most Enterobacteriales, in the Pseudoalteromonas and in most Aeromonadales (20). In most other bacterial species, there is no replication fork trap present, as experimentally demonstrated for the two circular *Vibrio cholerae* chromosomes (20). Thus, it appears the majority of bacterial species have little difficulty surviving without a replication fork trap. Indeed, a fork trap introduces a significant level of constraint to chromosome duplication: if one fork is stalled at an obstacle before it has progressed through the first five *ter* sites it encounters and cannot be reactivated, the second fork will be unable to rescue this blocked fork because it will be blocked by Tus-*ter* complexes. If the stalled fork cannot be reactivated, the cells are in danger of dying (3, 8), a problem that will not arise in the same way in bacterial chromosomes without a fork trap. We were recently able to recreate this scenario *in vivo*, by moving the origin from its original location into either the right-hand or left-hand replichore (41, 42). In these cells, replisomes coming from ectopic origins have to duplicate a significant proportion of the chromosome in an orientation opposite to normal, which results in replication-transcription conflicts that will delay the progression of one fork, whilst the second fork is stalled at Tus-*ter* complexes (8,41,42). The resulting cells show a significant growth defect, indicating that this scenario can cause serious issues for the cell. The observed growth defect is significantly alleviated if Tus is absent, demonstrating that preventing forks from being trapped is advantageous (8, 41, 42).

The results of this study highlight yet another constraint that is introduced by the replication fork trap: processing beyond that provided by standard replisome components is required when two replisomes converge at Tus-*ter*, otherwise the chromosome will remain under-replicated (Figure 3). While the majority of fork fusions take place away from Tus-*ter* complexes *in vivo* (16, 24), delays by one of the two replisomes are frequent enough to allow the detection of forks stalled at Tus-*ter* complexes (27). Thus, cells deal with this scenario on a regular basis. The fork trap must therefore provide a significant advantage to cells in order to compensate for the difficulties it can cause. The nature of this advantage is still unclear. Our previous results suggest that toxic intermediates arise in the termination area due to fork-fusion events. These intermediates are normally efficiently processed by proteins such as RecG helicase and 3’ exonuclease, but in their absence, lead to significant over-replication which is contained by the replication fork trap (3, 24). These intermediates can trigger cell lethality, leading to the speculation that the main purpose of the replication fork trap is to ensure that they only arise and are contained to a defined area of the chromosome where they can be efficiently processed (3). Cells without a termination area still have all the necessary proteins to process intermediates that might arise as a result of fork fusion events. However, acquiring the fork trap system has provided cells with the advantage of being able to confine the intermediates to a defined area of the chromosome, an advantage apparently strong enough to compensate for the constraints it introduces.

In conclusion, we have demonstrated that the fusion of two replication forks at Tus-*ter* results in incomplete DNA replication, leaving a gap of at least 15 bp of unreplicated DNA. Additional processing beyond that provided by established replisome components must be required for successful termination. The most obvious candidates, Rep, UvrD and RecG, are unable to promote replication to completion in this scenario, suggesting that another as yet unidentified molecule is likely to participate in replication termination at Tus-*ter*. Further investigation is required to understand the comprehensive requirements for completing DNA replication at Tus-*ter*, for understanding how and when Tus is displaced from the DNA during fusion and how the final stretch of DNA across the *ter* site is replicated. Such investigations will not only shed light on the molecular mechanics of the termination process, but also the precise reason why acquiring a replication fork trap is advantageous. Ultimately, they will lead to a better understanding of the factors that have shaped the overall landscape of bacterial chromosomes.

## ACKNOWLEDGEMENTS

The authors would like to thank Peter McGlynn and Ed Bolt for their critical reading of the manuscript. We would also like to thank Robert Lloyd and Geoff Briggs for kindly providing RecG.

